# Megabouts: a flexible pipeline for zebrafish locomotion analysis

**DOI:** 10.1101/2024.09.14.613078

**Authors:** Adrien Jouary, Pedro T.M. Silva, Alexandre Laborde, J. Miguel Mata, João C. Marques, Elena M. D. Collins, Randall T. Peterson, Christian K. Machens, Michael B. Orger

## Abstract

Accurate quantification of animal behavior is crucial for advancing neuroscience and for defining reliable physiological markers. We introduce Megabouts (megabouts.ai), a software package standardizing zebrafish larvae locomotion analysis across experimental setups. Its flexibility, achieved with a Transformer neural network, allows the classification of actions regardless of tracking methods or frame rates. We demonstrate Megabouts’ ability to quantify sensorimotor transformations and enhance sensitivity to drug-induced phenotypes through high-throughput, high-resolution behavioral analysis.

---

Advances in machine learning and computer vision have significantly improved the ability to track animal posture (Mathis and Mathis, 2020) and decompose postural time series into a sequence of distinct behavioral motifs (Wiltschko et al., 2015). These developments have driven significant progress in neuroscience (Pereira et al., 2020) and preclinical research (Wiltschko et al., 2020). Yet, comparing behavioral data across laboratories remains challenging due to differences in experimental setups, tracking methods, and analysis pipelines.

The larval zebrafish is well-suited for behavioral analysis, as its locomotion is naturally segmented into a burst-and-glide pattern. Previous research has employed unsupervised clustering methods to categorize these swim bouts and unravel the animal’s behavioral repertoire (Marques et al., 2018; Johnson et al., 2020; Mearns et al., 2020). However, these swim classification methods depend on specific camera resolutions, frame rates, tracking algorithms, and analysis pipelines, making it difficult to compare behavioral phenotypes across laboratories (von Ziegler et al., 2021). Megabouts tackles this challenge by providing a flexible computational pipeline to homogenize data from different sources (see Fig. S1). Its primary goal is to process zebrafish tracking time series data, segment it into individual swim bouts, and classify each bout according to the zebrafish’s locomotor repertoire (Fig. 1a,b). While focusing on freely swimming behavior, an add-on pipeline accommodates head-restrained assays which require specialized behavior segmentation and analysis.

**Figure 1.**
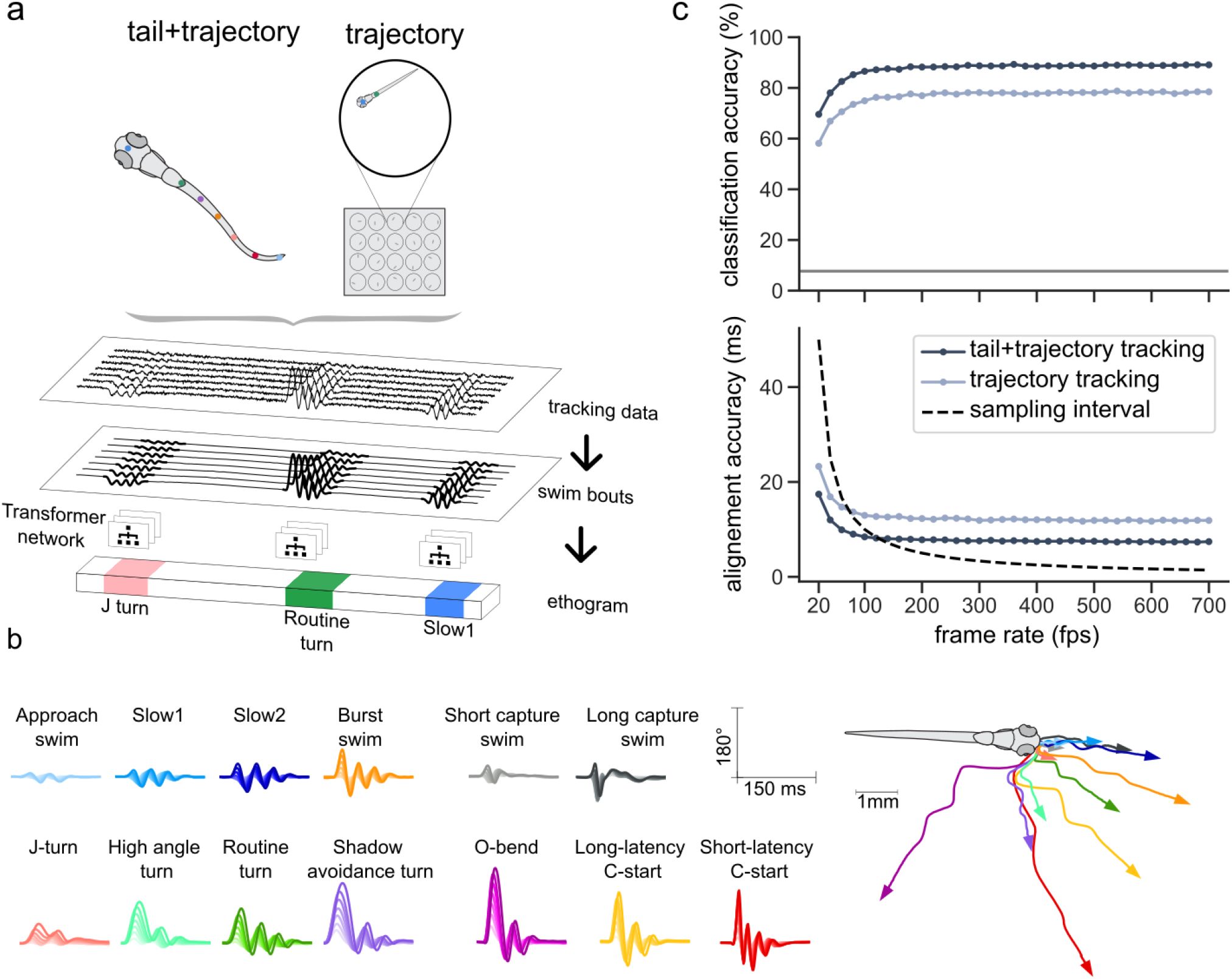
Megabouts pipeline overview and performance. **(a) Overview of Megabouts’ pipeline**. Megabouts can process either high-resolution tail and trajectory tracking or low-resolution trajectory tracking. The pipeline begins with data loading and preprocessing, followed by swim bout identification via a segmentation algorithm. A Transformer neural network then classifies each swim bout and predicts the location and sign of the first tail beat. **(b) Behavioral repertoire of zebrafish larvae composed of 13 swim bout categories**. Left: Exemplar tail angle time series for each category, representing bouts closest to the category mean. Right: Corresponding head trajectories with arrows indicating final head orientation. **(c) Performance of the Transformer neural network**. Top: Balanced classification accuracy at different frame rates. The downsampled input data was either presented with or without tail masking. Accuracy remains an order of magnitude above chance level (7.7%) in all conditions. Bottom: 95% confidence interval for the predicted location of the first tail beat. This first tail beat is predicted from the Transformer Neural Network along with the bout category. For frame rates below 100 fps, the prediction accuracy is below the data sampling interval, achieving super-resolution.

Zebrafish freely swimming behavior is typically studied in shallow arenas, recorded from above using video imaging. Tracking can be achieved with opensource software designed for zebrafish (Mirat et al., 2013; Štih et al., 2019; Guilbeault et al., 2021) or deep learning-based image processing (Mathis and Mathis, 2020; Pereira et al., 2022). However, there is considerable variation in the spatial and temporal resolution, with a trade-off between resolution and throughput. Megabouts accommodates these variations (see Fig. S2) by supporting two configurations for freely swimming behavior: ‘tail+trajectory tracking’, which involves high-speed tracking (100 to 2000 fps) of multiple tail landmarks, as well as head position and angle (cf. SV1); and ‘trajectory tracking’, which requires only the fish’s position and head angle, making it suitable for high-throughput assays (cf. SV2).

To validate Megabouts across different tracking configurations, we used a dataset of 4 million swim bouts recorded at high resolution (see Supp. Note 3.A). The segmentation and classification of these bouts (Fig. 1b) using the method from Marques et al., 2018 provided our reference labels. Using downsampling and masking strategies (Fig. S4c), we tested the software’s performance at lower frame rates and with trajectory-only tracking, ensuring high accuracy across conditions.

Segmenting tracking data into swim bouts is straightforward, given zebrafish’s burst-and-glide swimming pattern. We developed a segmentation algorithm that accurately (>94%, cf. Fig. S3 and Supp. Note 4.F.1) detects swim bouts across all frame rates using either tail oscillation speed or head positional and yaw velocities (Supp. Note 4.C).

Once the tracking data is segmented into swim bouts, we characterize the swim bouts using a Transformer neural network (Vaswani, 2017, cf Fig. S4b and Supp. Note 4.E). This architecture is well-suited for dealing with missing data. Each input token to the Transformer contains information about the larva’s posture and position at a single time step. The Transformer’s selfattention mechanism identifies relationships between posture measurements and focuses on the most relevant features. This contrasts with previous methods that rely on time series distance (Mearns et al., 2020 or feature extraction Marques et al., 2018, which are disrupted by missing data, temporal misalignment or frame rate variations. The Transformer architecture overcomes these limitations, offering robust classification across diverse conditions.

We trained the network using a supervised learning approach, the output consists of key movement features: bout category and subcategory, direction, and timing of the first tail beat. We found that the network’s performance reached a plateau at around 120 fps, and the classification accuracy remained high even without tail tracking (Fig. 1c, Fig. S5a). The first tail beat location was also predicted with high accuracy (Fig. 1c), achieving super-resolution for frame rates below 100 fps.

After validating Megabouts’ performance, we explored several use cases that highlight how its detailed behavioral descriptions improve upon coarser measures of behavior.

Understanding how sensory stimuli map to specific motor outputs is central to neuroscience. Megabouts’ swim bout categories were specifically designed to preserve these sensorimotor relationships (Marques et al., 2018). For example, while O-bend and Short Latency C-Start are both escape-like behaviors, they are known to be triggered by different stimuli (Burgess and Granato, 2007; Bhattacharyya et al., 2017). We found that Megabouts improved the mapping between external stimuli and the fish’s actions (Fig. 2a) as measured by the mutual information (see Supp. Note 4.F.2 and Fig. S6 for subcategory details). Bout direction extracted from the first tail beat laterality is also significant. For instance, escapes are more likely generated contralaterally during aversive stimuli like looming or approaching dot (Fig. 2a). Interestingly, forward movements such as Slow 2 are also directionally biased, occurring more ipsilateral to the grating direction (Fig. 2a). These results illustrate the advantage of using a detailed library to accurately quantify the ‘tuning curve’ of actions.

**Figure 2.**
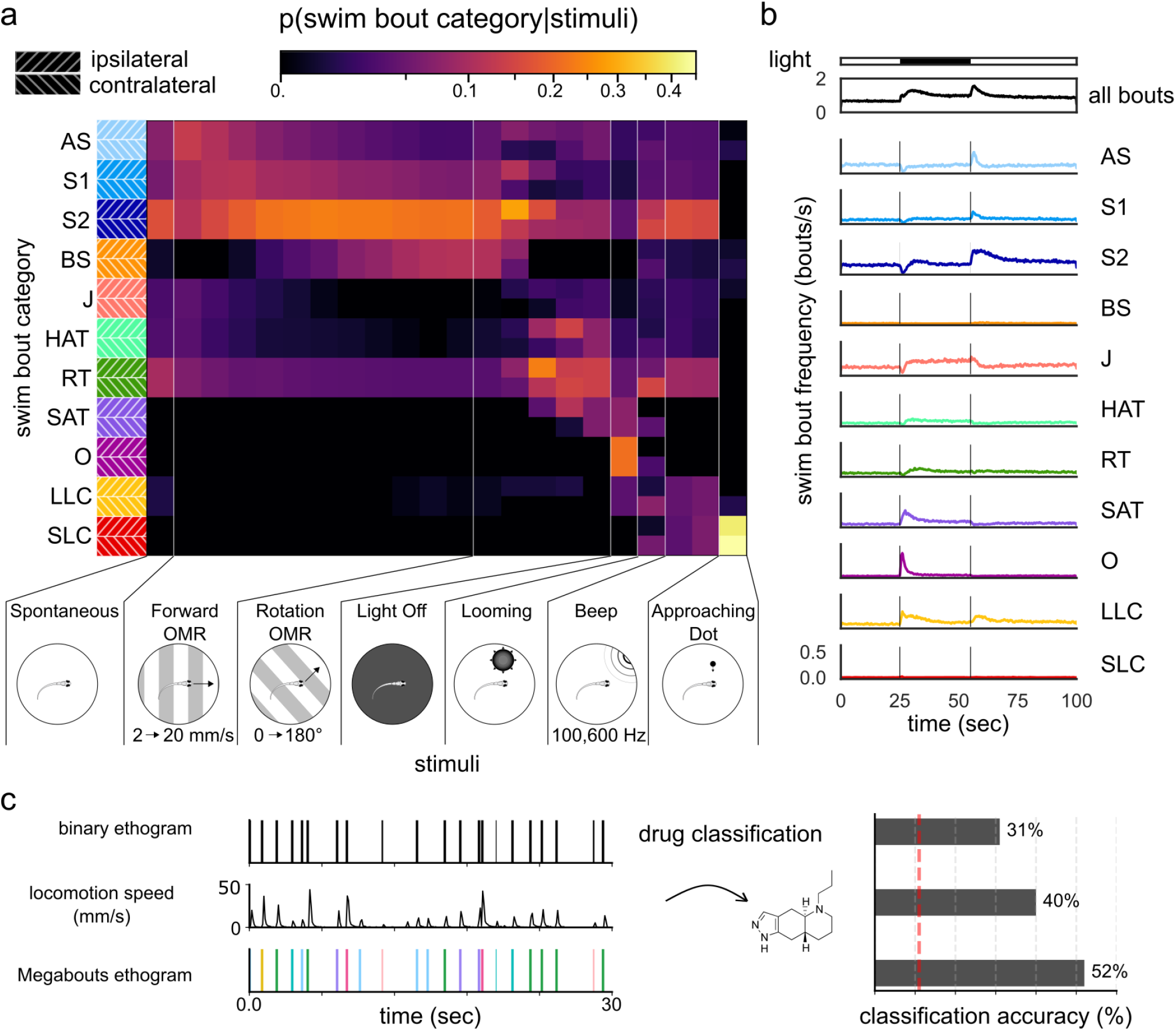
Sensorimotor coupling and behavioral phenotyping. **(a) Sensorimotor coupling**. Conditional probability of swim bout occurrence given a stimulus, computed from 1.95 million bouts from 108 larvae. Bout categories are color-coded as in Fig. 1b. For directional stimuli, bouts are split into ipsilateral or contralateral based on the direction of the first tail beat relative to the stimulus. **(b) Temporal dynamics of swim bouts recorded using Zebrabox**. Average frequency of swim bout initiation for each category as a function of time relative to the light on/off stimuli, averaged over 4 trials for N=372 larvae. The subplots for each category share the same y-scale. **(c) Enhanced phenotyping of neuroactive drugs**. MiniRocket classifier trained on three behavioural time series: binary movement (top), locomotion speed (middle) and Megabouts action categories (bottom). Vertical dotted line is chance level at 11%.

Megabouts can also uncover the temporal structure of behavior. The light on / off protocol is popular for high-throughput phenotyping (Kokel et al., 2010; Rihel et al., 2010). Typically, behavior quantification is performed using measures of the animal’s overall locomotion, such as the frequency of swim bouts, modulated by light depending on the fish’s condition (Fig. 2b). Using Megabouts on data from a commercial recording system at 25 fps, we broke down the locomotion time series into temporal profiles with distinct time scales and on/off selectivity (Fig. 2b). Notably, we observed consistent O-Bend responses after light off across different well sizes (see Fig. S7 and Fig. SV3). This allows for analyses that were previously limited to high frame rate cameras (Randlett et al., 2019) and opens possibilities to analyze action sequences to detect latent internal states and long-term modulation (Reddy et al., 2022).

Finally, we asked whether the rich action-level description provided by Megabouts could sharpen the behavioural fingerprint of drugs, building on work showing detailed behavioral quantification in mice improved phenotyping(Wiltschko et al., 2020). We recorded 162 larvae exposed to nine neuroactive compounds while presenting a battery of visual stimulus (see Supp. Note 2.B). We then trained a MiniRocket time-series classifier citepdempster2021minirocket to identify the compound from behavioural data represented at three different resolutions: a binary “movement vs. no-movement” trace; the head locomotion speed; or the full Megabouts action-category time series. Prediction accuracy rose from 31% with the binary trace to 40% with speed, and leapt to 52% with action categories (chance=11%; Fig. 2c). Thus, fine-grained segmentation of behaviour substantially boosts the sensitivity and specificity of phenotype-based screening.

Megabouts also includes a pipeline for head-restrained experiments (see Supp. Note 5), commonly used for neural imaging. Head fixation leads to postures not seen during freely swimming, variable duration and modulation of frequency within a single movement (Portugues and Engert, 2011), making free-swimming classification methods unsuitable. To quantify the less stereotyped, variable-length tail movements, we developed a new approach based on convolutional sparse coding (Wohlberg, 2017; Cogliati et al., 2017). This method is grounded in the hypothesis that complex tail movements are generated by sparse control signals driving different tail oscillation motifs (Mullen et al., 2024). Convolutional sparse coding represents the tail angle as a combination of motifs, learned from the data: slow, fast, turn, and escape/struggle (see Fig. S8a).

We validated this approach using optomotor behavior, where the fast motif was progressively recruited with increasing grating speed (see Fig. S8c,d), consistent with free-swimming observations (Severi et al., 2014). These motifs provide refined motor regressors for neural activity analysis.”

Finally, the website megabouts.ai offers extensive documentation on the code base and provides tutorials for common use cases. By providing a common atlas for zebrafish behavior, the Megabouts Python package has the potential to foster cross-disciplinary collaborations between neuroscience, genomics, and pharmacology, accelerating scientific discovery.

## Code availability

Megabouts is available under a non-commercial research and academic license. All software, algorithms and tools are available on GitHub: at https://github.com/orger-lab/megabouts. The full documentation of Megabouts is available at megabouts.ai.

## Acknowledgments

We thank Emily G. Tippetts for her guidance on the Zebrabox setup, and Dean Rance and Carolina Gonçalves for their valuable insights on the Transformer architecture. Special thanks to Leandro Scholz for his feedback on the manuscript and for being an early user of the package. We also appreciate the helpful feedback from other early users: Claudia Feierstein, Joaquim Contradanças, Katharina Kötter, Daguang Li, and Emiliano Marachlian. Lastly, we are grateful to the Champalimaud Research Fish Platform for their logistical support. This work was realized through funding to M.B.O from an ERC Consolidator Grant (ERC NEUROFISH 773012) and to A.J. from the La Caixa Foundation (LCF/BQ/PR20/11770007). C.K.M and A.J. received support from Fundação para a Ciência e Tecnologia (FCT-PTDC/BIA-OUT/32077/2017-IC&DT-LISBOA-01-0145-FEDER). J.C.M. received the support of a fellowship from “la Caixa” Foundation (ID100010434, LCF/BQ/PR21/11840005). E.M.D.C. received support from Fundação para a Ciência e Tecnologia (Bolsa SFRH/BD/147089/2019). M.B.O and C.K.M received support from the Champalimaud Foundation; C.K.M from the Simons Collaboration on the Global Brain (543009); and M.B.O from the Volkswagen Stiftung Life? Initiative. We received support from the research infrastructure CONGENTO, co-financed by the Lisboa Regional Operational Programme (Lisboa2020) under the PORTUGAL 2020 Partnership Agreement, through the European Regional Development Fund (ERDF) and Fundação para a Ciência e Tecnologia (Portugal), under the project LISBOA-01-0145-FEDER-022170.

## Author contributions

A.J. and P.T.S. designed and performed high-resolution freely swimming behavioral experiments. J.C.M. developed the prey-capture assay and collected the corresponding dataset. E.M.D.C. developed the head-restrained assay and collected the corresponding dataset. J.M.M. and A.J performed pharmacological treatment experiments and collected data. A.J. and R.T.P. designed and conducted Zebrabox experiments. A.L., A.J., and M.B.O. developed the tracking software and visual stimuli presentation code, and contributed to the design of the analysis pipeline. A.J. performed the data analysis and developed the Megabouts software. M.B.O. and C.K.M. contributed to the design of the analysis pipeline and supervised the research. All authors contributed to the manuscript.

## Supplementary Figure

**Figure S1.**
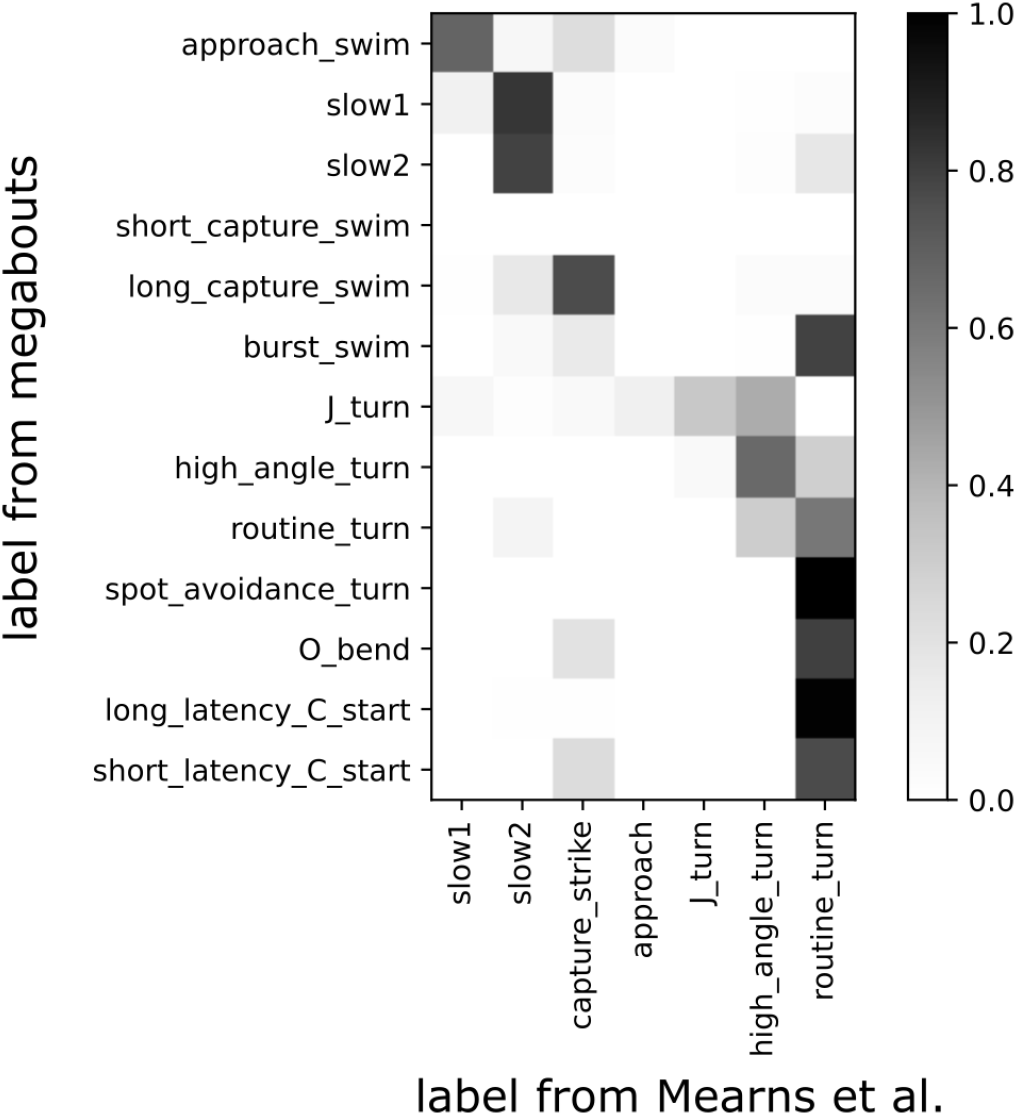
Comparison of Megabouts Bout Classification with Labels from Mearns et al., **2020**. This contingency table compares swim bout categories classified by Megabouts (rows) and those from Mearns et al., 2020 (columns), computed from approximately 40,000 bouts in the Mearns et al., 2020 dataset. Each cell shows the proportion of bouts in a Megabouts category that match a Mearns et al., 2020 category, normalized by the total number of bouts in each Megabouts category (values from 0 to 1). Darker shades indicate higher correspondence. The differences, even for similarly named categories, highlight the need for a standardized classification provided by Megabouts.

**Figure S2.**
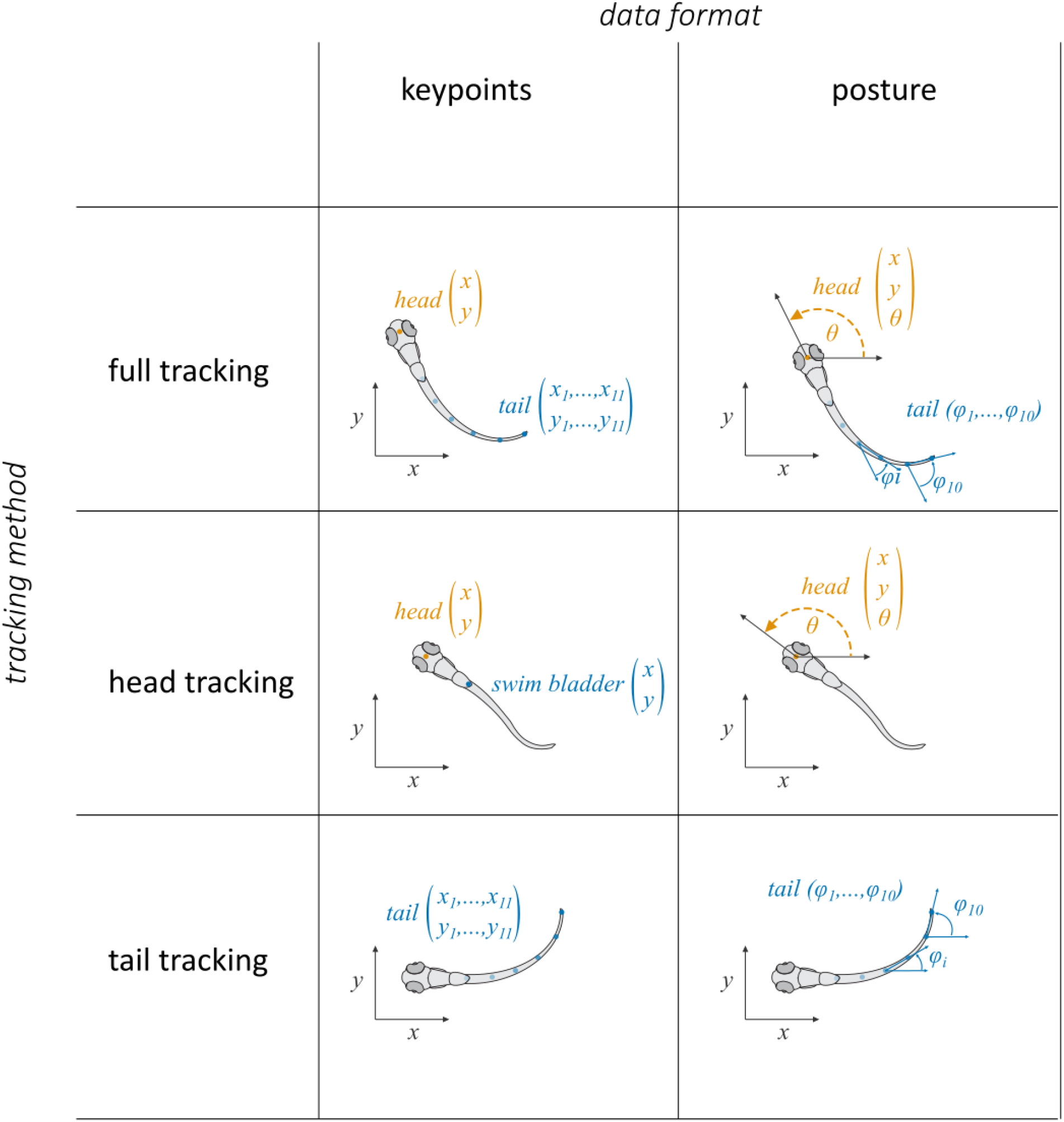
Tracking configuration compatible with Megabouts. Megabouts supports multiple tracking configurations: ‘tail tracking’ (for head-restrained conditions), ‘head tracking’, or ‘full tracking’. Head tracking requires two keypoints to estimate the position and orientation of the larva. Tail tracking requires at least four keypoints from the swim bladder to the tail tip. Data can be provided as keypoint coordinates or as posture variables, including head yaw and tail curvature.

**Figure S3.**
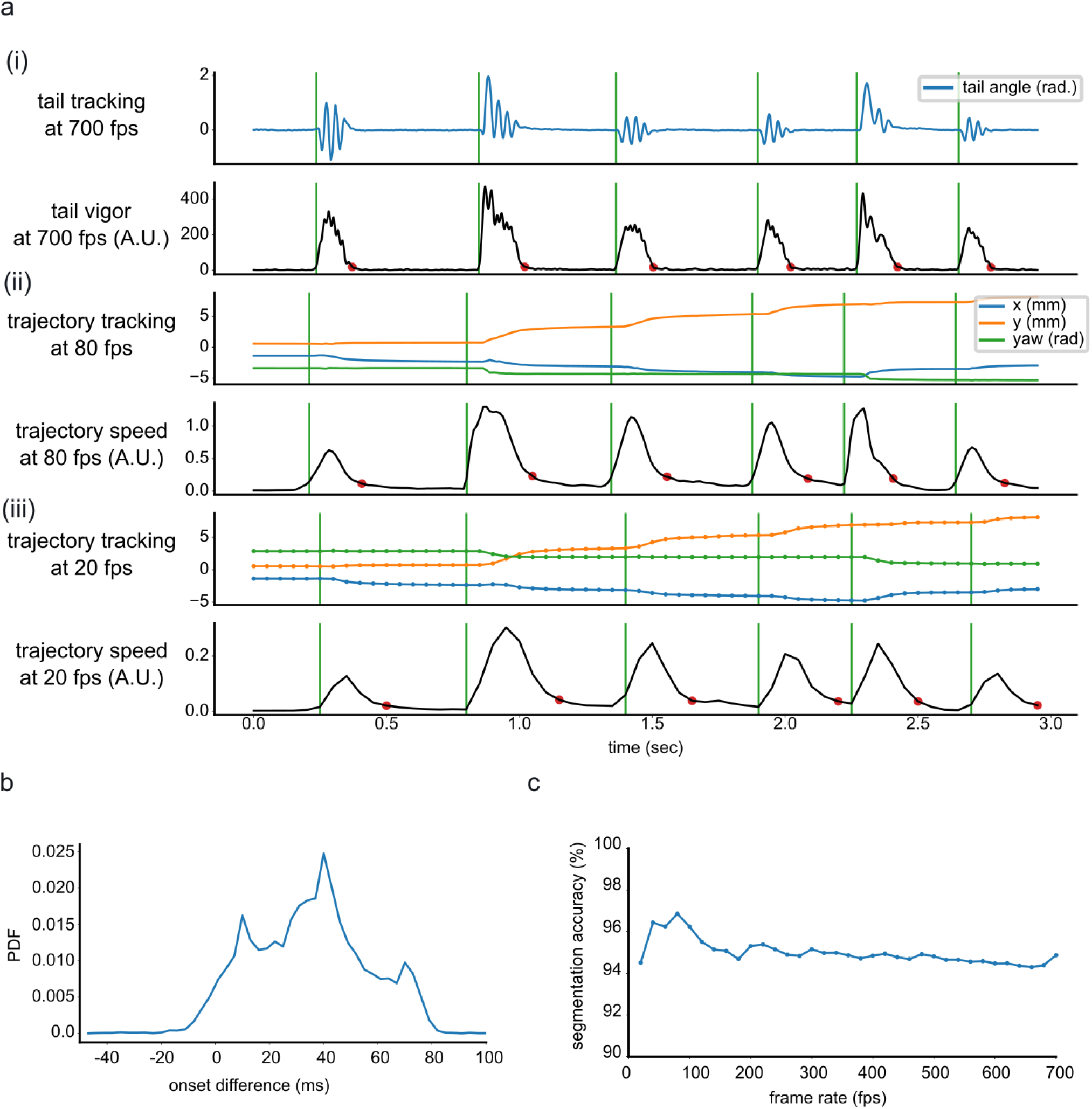
Segmentation algorithm and performance. **(a) Illustration of the segmentation algorithm**. **(i) High frame-rate segmentation**. A 10-second sample of tail tracking at 700 fps is shown. By applying a threshold to the tail vigor, we can identify the onset (green line) and offset (red dot) of swim bouts. **(ii) Downsampled to 80 fps with tail masking**. The same data is downsampled to 80 fps. Trajectory speed is computed based on the derivative of x,y and yaw coordinates. First, a peak-finding algorithm is used to locate the maximum trajectory speed. Then, a threshold of 20% of this peak value is applied to determine the onset and offset of swim bouts. **(iii) Downsampled to 20 fps with tail masking**. The same process as in (ii) but with downsampling to 20 fps. **(b) Onset jitter**. The distribution of the difference between onset times computed using tail angle vigor at 700 fps and trajectory at 20 fps is shown. The analysis is based on N=4 larvae from high-resolution recordings. **(c) Segmentation accuracy**. A reference segmentation was computed from tail angle at 700fps using 20 larvae. This reference was compared with segmentation obtained from trajectory-only data downsampled from 20 to 700 fps. See Supp. Note 4.F.1.

**Figure S4.**
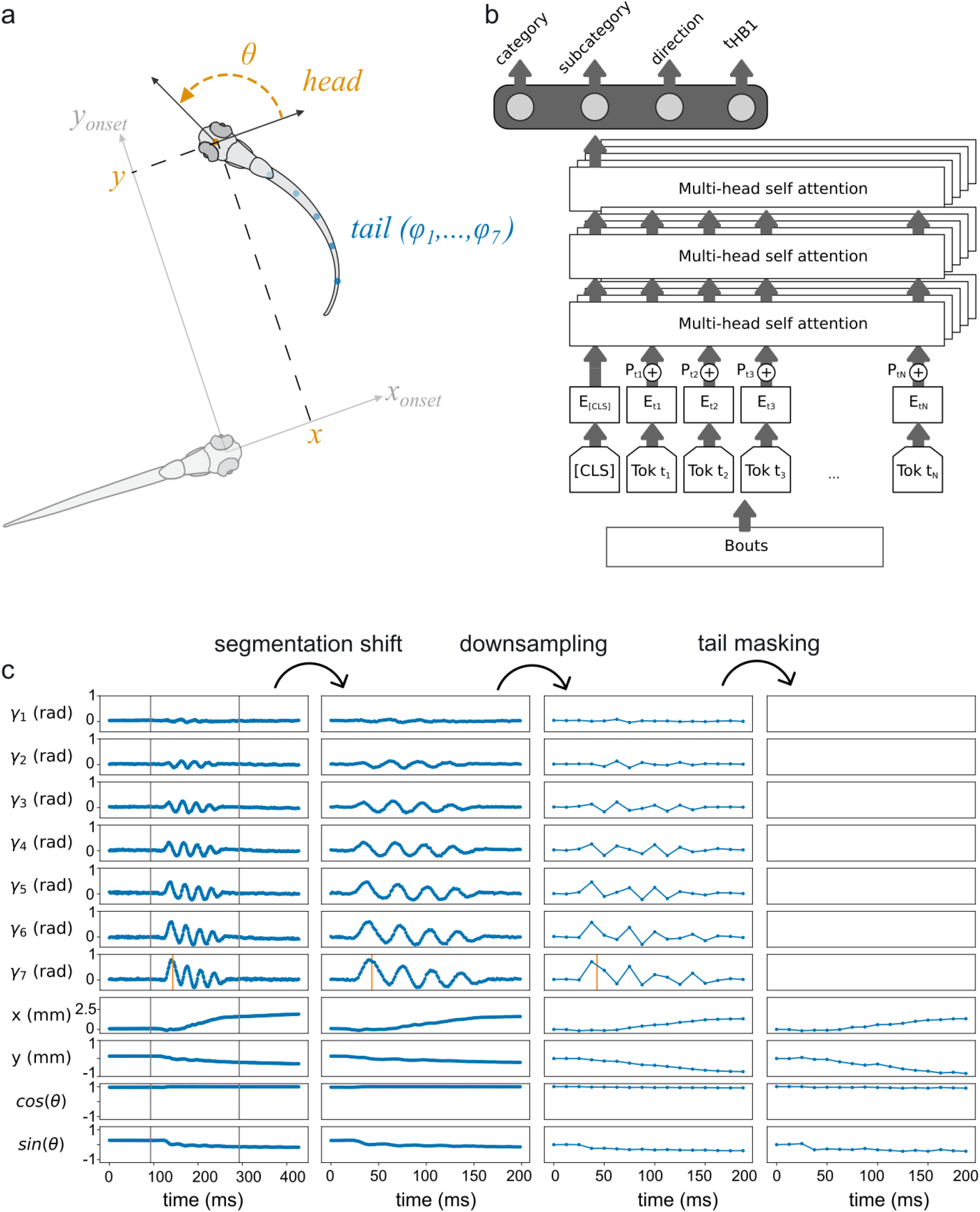
Transformer input and architecture. **(a) Input data for the Transformer**. For each swim bout, the first seven out of ten tail angles (*γ*_1_, …, *γ*_7_) are used as input, along with coordinates computed after the bout onset, relative to the fish’s position and orientation at onset. **(b) Transformer architecture**. Each token [*T ok t*_*i*_] corresponds to the pose measured at time *t*_*i*_ relative to the movement onset. [*T ok t*_*i*_] is a vector containing 11 values: tail angles (*γ*_1_(*t*_*i*_),…, *γ*_7_(*t*_*i*_)), *x*(*t*_*i*_) and *y*(*t*_*i*_) coordinates, and the head angle cos(*θ*(*t*_*i*_)), sin(*θ*(*t*_*i*_)). Missing measurements are replaced with a fixed learned value. After passing through an embedding layer, temporal embeddings are added. The tokens are then processed by 3 layers of 8-head multi-head self-attention. Finally, outputs are computed from the learned classification token [CLS]. **(c) Illustration of downsampling/masking data augmentation**. The process begins with an example bout. Each line corresponds to a token dimension. A random shift is applied to the segmentation (two vertical black lines) to achieve translational invariance in bout classification and jitter the first tail beat location (shown as the vertical orange line). The time series is then downsampled, and masking can omit the tail information.

**Figure S5.**
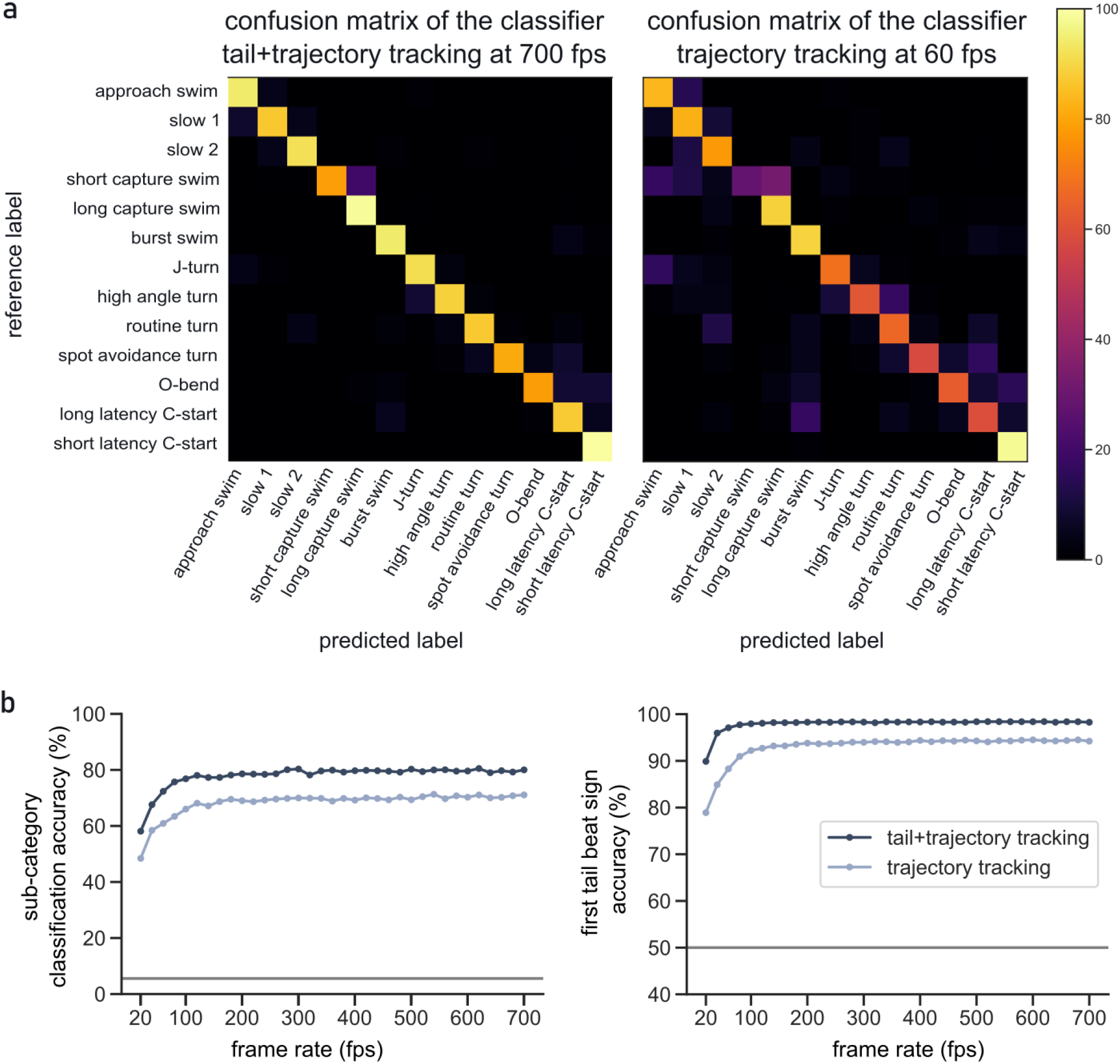
Classification accuracy and bout directionality. **(a) Confusion matrix**. Left: Confusion matrix at high resolution, corresponding to a balanced accuracy of 89.1%. Right: Confusion matrix using only trajectory data downsampled to 60 fps, corresponding to a balanced accuracy of 71.2%. **(b) Performance in bout subcategory and directionality**. Left: Balanced classification accuracy for swim bout subcategories across different frame rates. The downsampled input data were presented with or without tail masking. Right: Accuracy of the estimation of the first tail half beat amplitude sign, which determines the directionality of the movements.

**Figure S6.**
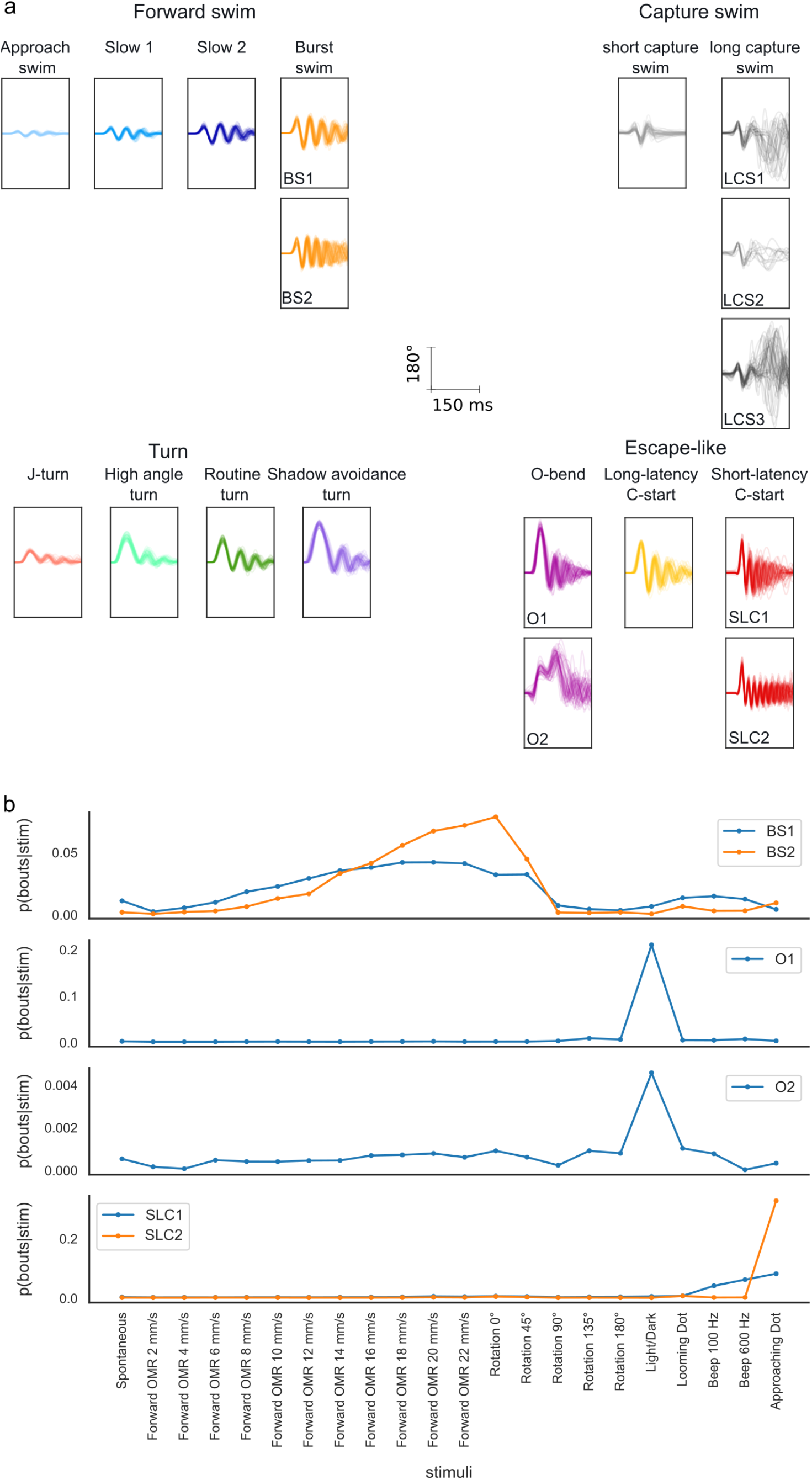
Swim bouts by subcategory. (**a**) For each swim bout subcategory, 50 swim bouts were selected from those in the test dataset that were classified with high probability (>98%). (**b**) We illustrate the probability of bouts occurring from specific subcategories given the stimulus presented. The O-bend subcategories O1 and O2 are presented in different plots since they have very different probabilities of occurrence.

**Figure S7.**
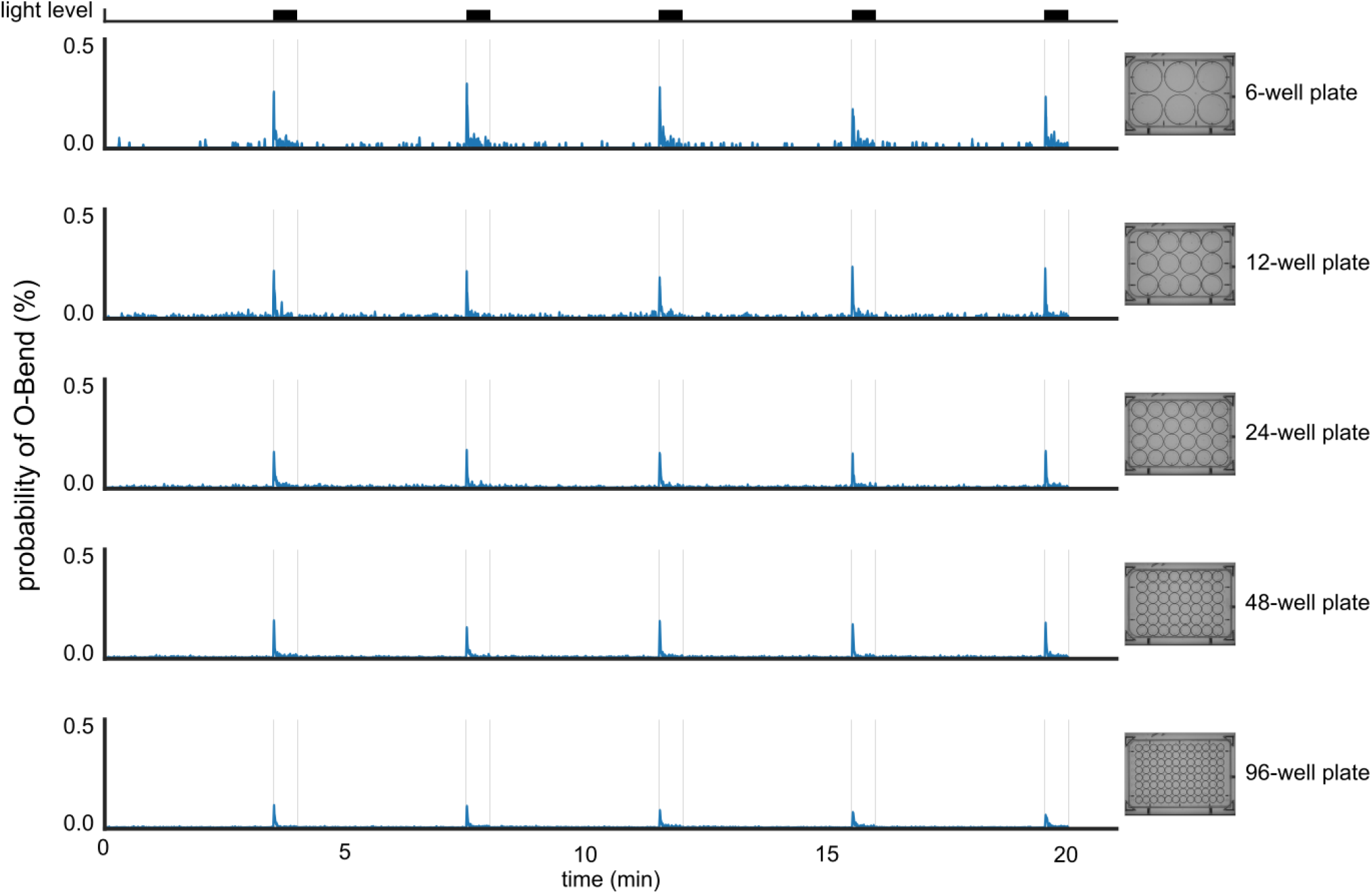
Occurrence of O-Bend in response to light stimuli. Probability of occurrence of O-Bend across time for different well sizes. The probability was computed using N=2 wells for each size. The probability was smoothed using a 1-sec box-car filter.

**Figure S8.**
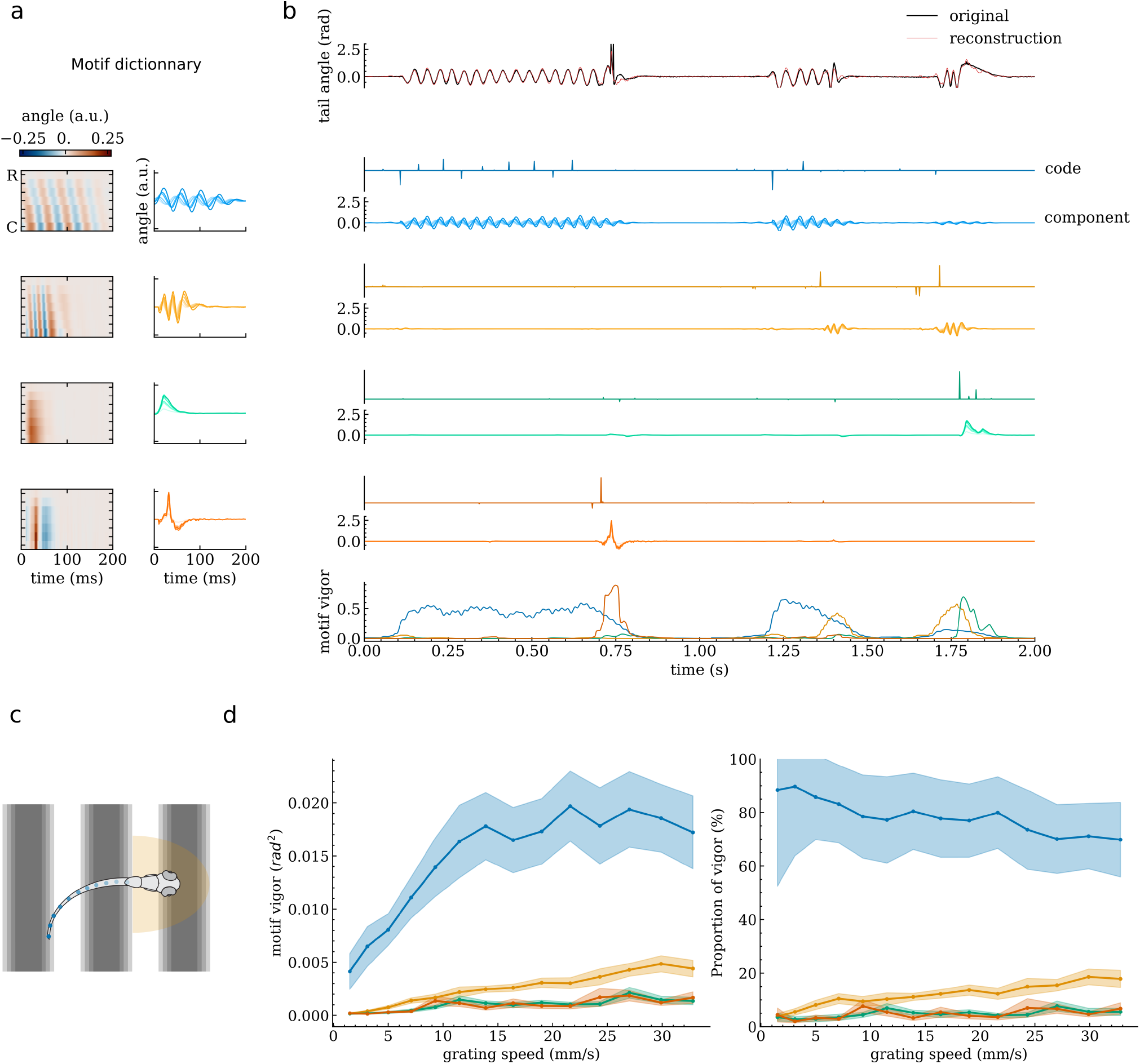
Head-restrained pipeline for motif analysis. **(a)** Illustration of the four motifs learnt from the data: slow, fast, turn, struggle. Left: heat map of each motif shows tail beat propagation along rostral-caudal segments and time. Right: Same data as time series with increasing contrast along rostral-caudal axis. (The motif color code is shared in a., b. and d.). **(b)** Illustration of sparse coding and motif vigor. Top. The original tail angle (black) is reconstructed by adding the four components (red). Middle. Each component is computed by the convolution of a sparse code and the motif. Bottom. The contribution of each motif can be computed as the rolling variance of each component (we used a window of 40ms) **(c)** A grating moving at different speeds was displayed below a head-restrained larva. A virtual-reality system was used to adjust the speed of the moving grating depending on the fish tail speed. **(d)** For each motif we computed the vigor (left) or proportion of motif recruitment (right) as a function of the grating speed (filled area corresponds to mean±std.error, N = 25 larvae).

## Supplementary Information

### Supplementary Note 1: Fish Care

Adult fish were maintained at 28^*°*^C on a 14:10 hour light cycle. Embryos were collected and larvae were raised at 28^*°*^C in E3 embryo medium (5 mM NaCl, 0.17 mM KCl, 0.33 mM CaCl2 and 0.33 mM MgSO4) in groups of 25. Sexual differentiation occurs at a later stage, and therefore the sex of the animals cannot be reported. Behavioral experiments were conducted using the wild-type line Tubingen (Tu) between 5 and 7 days post fertilization (dpf). Larvae were fed with rotifers (*Brachionus sp*.) from 5 dpf onwards. All experimental procedures were approved by the Champalimaud Foundation Ethics Committee and the Portuguese Direcção Geral Veterinária, and were performed according to the European Directive 2010/63/EU. The Zebrabox experiments were preformed according to the University of Utah’s Institutional Animal Care and Use Committee (IACUC) guidelines under protocol 00,001,487.

### Supplementary Note 2: Behavioral experiments

#### 2.A. High-resolution experiments

##### 2.A.1. Setup

The behavior of each larva was recorded in a circular acrylic arena carved and polished using a CNC machine (Núcleo de Oficinas, Instituto Superior Técnico). The arena had a diameter of 50 mm and a depth of 4 mm, with rounded edges achieved by a progressive bevel with a radius of curvature of 11 mm. Fish were imaged using an infrared array consisting of 64 LEDs at 850 nm (TSHG6400, Vishay Semiconductor) controlled by a T-Cube LED Driver (LEDD1B, Thorlabs), a fixed focal length lens (86-207, f = 16 mm, Edmunds Optics), an infrared long-pass filter (LP780-37, 780 nm, VisionLightTech), and a high-speed camera (EoSensCL MC1362, Mikrotron). Camera images were acquired using a frame grabber (PCIe-1433, National Instrument). The camera settings allow to image a region of 948*948 pixels at 700 fps and a shutter time of 1.423 ms. For visual stimulation, we used a video projector (ML750e, Optoma) along with a neutral density filter (NE06A, Thorlabs) and a cold mirror (64-452, Edmunds Optics) to project an image onto a diffuser screen (three layers of Rosco Cinegel White Diffuser 3000) positioned 5 mm below the larva. A transducer speaker (4501, VISATON) glued to the acrylic plate beneath the fish area delivered acoustic stimuli.

##### 2.A.2. Behavioral tracking

The posture was computed in real-time using a custom-written C# software. This software tracks both eyes and follows the tail curve using 10 segments, starting at the swim bladder.

Image processing is performed on background-subtracted images. The background for each pixel was computed using the mode of the distribution of pixel values recorded in 100 frames sampled over a few seconds to ensure sufficient fish movement. Using the mode rather than the mean prevents contamination of the background by the fish image, assuming that each pixel is imaged in absence of the fish most of the time. A large Gaussian kernel is applied to smooth the background-subtracted image. The maximum value indicates the fish location. From this maximum, a flood-fill algorithm is employed to obtain a binarized image of the fish trunk. The principal axis of the fish trunk is computed from the moments of the binarized region. The fish head direction is then determined by finding the point along the principal axis where the total intensity along the orthogonal line is maximum. This location corresponds to the mid-eye. The relative position of the eye to the fish centroid provides a robust identification of the fish heading direction.

From this, the tail is found by following an iterative arc-search algorithm. This algorithm tracks the skeleton of the fish by localizing the next keypoints along an arc of ± 60^*°*^ around the direction from the previous keypoints. Keypoints are found by computing the weighted mean along the arc (with the weight being zero below a given threshold or equal to the pixel value of the background-subtracted image).

The algorithm runs in real-time at 700 Hz. It leverages the fast frame rate by limiting the search to a region of interest around the fish position in the previous frame. Time consuming full-frame operations are avoided; for instance, thresholding is performed only on pixels along the arc rather than on the entire image.

##### 2.A.3. Visual and acoustic stimuli

We use the term ‘virtual open loop’ to describe a configuration where, although an experimental closed-loop system presents visual stimuli relative to the fish’s position and orientation, the fish’s displacement does not affect the stimuli’s relative position, effectively creating an open-loop condition.

The stimuli presented consisted of:

- **Approaching Dot**. This stimulus is displayed in a virtual open loop. A black disk (1 mm radius), initially positioned 2 cm away from the fish, approaches at a speed of 5 mm/s. After the approach, the disk remains below the fish for 1 second. The disk moves in a direction that is perpendicular to the fish’s head vector (± 90^*°*^).
- **Directional Optomotor Response**. This stimulus is displayed in a virtual open loop. A moving grating with a spatial period of 10 mm moves at 10 mm/s at an angle of either 0^*°*^, 45^*°*^, 90^*°*^, 135^*°*^, 180^*°*^, 225^*°*^, 270^*°*^, or 315^*°*^ relative to the fish’s heading. For each direction, the grating is first presented at a speed of 0 below the fish for 5 seconds, then moves at 10 mm/s for 10 seconds.
- **Looming**. This stimulus is displayed in a virtual open loop. A black disk, with its center 4 cm away from the larva, increases its radius at a linear speed of 10 mm/s until it reaches the larva after 4 seconds, at which point it disappears. The disk center is located in a direction that is perpendicular to the fish’s head vector (±90^*°*^).
- **Acoustic Startle**. A 100 ms pulse at either 100 Hz or 600 Hz. The sensorimotor mapping displayed in Fig. 2a counts the bout category probability up to 2 seconds following the pulse onset.
- **Forward Optomotor Response**. A moving grating with a spatial period of 10 mm and a velocity of 8, 16, or 24 mm/s is displayed along the fish axis. This stimulus orientation was in an open loop; regardless of how much the fish rotated, it was always aligned with the grating. However, a closed-loop was used for forward locomotion. The phase of the grid was computed as a function of the forward speed of the fish. Thus, if the fish swims at a speed matching the grating, it would see a static grating orthogonal to its swimming direction.
- **Light/Dark**. Switch the projector intensity from 1000 lux to 0 for 30 seconds. The sensorimotor mapping computed in Fig. 2a counts the bout in the first 4 seconds following the switch.

Visual stimuli were displayed at 60 fps using a custom-written rendering engine using OpenTK and generated using OpenGL Shaders. After 8 minutes of habituation, stimuli were presented in 6 pseudo-random order blocks, including inter-stimuli intervals of 3 minutes. The experiment lasted 3 hours.

##### 2.A.4. Prey Capture

Prey capture assays were performed as described previously (Marques et al., 2018). The behavioral setup used in these experiments is similar to the one used for the other behaviors, but was fitted with a Schneider apo-Xenoplan 2.0/35 lens (Jos. Schneider Optische Werke GmbH, Germany) and had a higher spatial resolution (365 pixels per cm) to ensure correct tracking of the eyes and prey. Experiments used a 25 mm x 25 mm with 3 mm depth square arena and larvae were illuminated from above with white light (1000 lux). Fish were fed with 50-100 paramecia (*Paramecium caudatum*) or rotifers (*Brachionus plicatilis*) and were allowed to hunt for 1 to 2h. For a subset of the long capture swims larvae roll their body generating tail tracking mistakes. These bouts were excluded from the analysis.

#### 2.B. Pharmacological screening experiments

##### 2.B.1. Setup

We recorded larvae swimming in an array of 20 wells. Each larva was placed in an individual circular acrylic arena carved and polished using a CNC machine (Núcleo de Oficinas, Instituto Superior Técnico, Portugal). The arena had a diameter of 44 mm and a depth of 4 mm, with tapered edges achieved by a bevel slope of 3 mm, resulting in a bottom diameter of 38 mm. Fish were imaged using an EoSens 4CXP Camera (MC4086, Mikrotron) running at 400 fps. The frame grabber was a Silicon Software GmbH AQ8 CXP6D. The camera settings allow to image a region of 2336*1728 pixels at 400 fps, the field of view of the camera was 20 cm x 25 cm. Fish were illuminated by an infrared array consisting of 64 LEDs at 850 nm (TSHG6400, Vishay Semiconductor) controlled by a T-Cube LED Driver (LEDD1B, Thorlabs) with a spacing sufficient to cover the full FOV, a fixed focal length lens (Xenoplan 2.0/28, Schneider Optische Werke), an infrared long-pass filter (LP780-37, 780 nm,VisionLightTech). For visual stimulation, a video projector (ML750e, Optoma) and a cold mirror (64-452, Edmunds Optics) projected an image onto a diffuser screen (three layers of Rosco Cinegel White Diffuser #3000) positioned 5 mm below the larva.

##### 2.B.2. Behavioral tracking

Due to the lower resolution resulting from the large field of view (FOV), we limited the tracking to the larva’s position and orientation. The background subtraction and body tracking methods are similar to those described in Supp. Note 2.A, but they do not include the arc search for tail segments.

##### 2.B.3. Visual stimuli

The stimuli presented consisted of **Approaching Dot, Directional Optomotor Response** and **Light/Dark** described in Supp. Note 2.A. Note that this experimental protocol presented stimuli with a lower inter stimuli interval than in Supp. Note 2.A, reaching as little as 15 s between two approaching dot stimuli presentations. This was used to probe potential differences in habituation or fatigue

***Drug treatment***. Fish were treated with nine neuroactive drugs:

- Ketanserin (serotonin 5-HT2 antagonist, 8.5 µM)
- Quipazine (serotonin 5-HT2*/*3 agonist, 25 µM)
- Trazodone (serotonin antagonist and reuptake inhibitor, 25 µM)
- Quinpirole (selective D_2_/D_3_ receptor agonist, 25 µM)
- Clozapine (atypical antipsychotic, 8.5 µM)
- Fluoxetine (selective serotonin reuptake inhibitor (SSRI), 8.5 µM)
- Haloperidol (D_2_ receptor antagonist, 8.5 µM)
- Apomorphine (non-selective dopamine agonist, 25 µM)
- Valproic acid (GABAergic voltage-gated sodium channel blocker, 25 µM)

Stock solutions were prepared by dissolving the compounds in autoclaved Milli-Q water. For drugs with low water solubility, an initial solution in 10% (v/v) DMSO was prepared, ensuring that the final DMSO concentration in the stock did not exceed 0.3%, a threshold below which no behavioral changes have been reported. Drug stocks were stored at -20^*°*^C after confirming stability via UV-Vis spectroscopy.

To prepare the arena solution, the appropriate stock volume was diluted in 600 mM Tris-buffered E3 medium to a final volume of 5 mL. All arena solutions had a final pH of 7 ± 0.3. The concentration for each drug was set to half of the maximum non-lethal dose. Larvae were allowed to habituate to the arena and drug solution for 1 hour before data collection. Each drug and concentration was tested on 18 fish.

###### Dru. Classification Analysis

We applied the Mini-Rocket classifier to three different time-series: binary movement indicators, locomotion speed, and a multivariate one-hot encoded time series of movement categories obtained from Megabouts. We employed a 10-fold stratified cross-validation. Each dataset was transformed using MiniRocketMultivariate with 5000 kernels. The transformed data was then classified using a ridge classifier.

#### 2.C. Zebrabox experiments

##### 2.C.1. Setup

We used a commercial system for recording zebrafish (Zebrabox, Viewpoint, France). Each experiment lasted 20 minutes, and the light on/off stimulus sequence can be seen in Fig. S7. We recorded multi-well plates with 6, 12, 24, 48, and 96 wells. For each well size, we ran two experiments with different larvae. Each experiment produced a compressed movie with a resolution of 9 pixels per mm. This setup produces a good edge case, however, we encourage users to work with higher resolution or frame rate if possible. We found that the biggest limitation of the Zebrabox was the long exposure time resulting in very blurry images when the fish is swimming fast. The users should aim to increase the light intensity and decrease the exposure to get optimal results even at low frame rate.

##### 2.C.2. Behavioral tracking

To track the video obtained from the Zebrabox system, we used SLEAP (Pereira et al., 2022). We tracked two keypoints: one placed between the fish’s eye and another on the swim bladder. Each multiwell video was cropped into several single-well recordings. For each recording we computed the background using the mode of the pixel value distribution sampled along the recording. From the grayscale videos, we generated a composite color that showed the background and a background-substracted image in different color channels. Given the low-resolution, we found that using this composite color to underline the foreground was critical to help SLEAP deal with cases where the fish overlapped with the edge of the arena. SLEAP was trained using the top-down pipeline after labeling 1900 frames. Since each cropped video contained a single fish, we did not need to use cross-frame identity tracking.

#### 2.D. Head-restrained experiments

##### 2.D.1. Setup

Experiments were conducted in a cylindrical acrylic arena carved and polished using a CNC machine (Núcleo de Oficinas, Instituto Superior Técnico, Portugal) The larvae were mounted on a flattened cone made from Sylgard (SYLGARD 184 Silicone Elastomer Kit, The Dow Chemical Company). Each larva was recorded using a high-speed camera (EoSensCL 1362, Mikrotron) fitted with an infrared long-pass filter (LP780-37, 780 nm, VisionLightTech) to block visible light and a lens (M30.5mm, Edmund Optics) to optimally focus on it. The camera was positioned above the arena, which was illuminated from below using a mounted high-power infrared LED (M780L2, Thorlabs) and controlled by a T-Cube LED Driver (LEDD1B, Thorlabs). Visual stimuli were displayed using a LED projector (ML500, Optoma). Three cold mirrors (one below, two laterally) were placed around the arena for 270° visual stimulation. Behavior was recorded at 700 fps. Larvae were embedded the evening prior to recording using low-melting point agarose (UltraPure LMP Agarose, Cat#16520100, Invitrogen by Life Technologie). Their tails were freed by removing agarose in a square caudal to the swim bladder to allow for maximum tail movement. The arena was filled with E3, and larvae were allowed to recover and habituate overnight at 28^*°*^C.

##### 2.D.2. Tracking

The posture was computed in real-time using a custom-written C# software. The tail was tracked using the iterative arc-search described in Supp. Note 3.A.

##### 2.D.3. Stimuli

Forward gratings at a range of speeds (15 speeds, 0-32.8 mm/s) were displayed for 20 seconds in a randomised order, with 5 repetitions each. Gratings were stationary for 10 seconds between trials. The width of the stripes was adjusted to match the free-swimming stimuli (1 cm width at 5 mm distance). Grating speed was updated continuously using a closed-loop virtual reality system.

### Supplementary Note 3: Megabouts training dataset

#### 3.A. Freely swimming

To train the Transformer neural network, we gathered 1.95 million swim bouts from 108 larvae from the experiments described in 2.1. We merged this dataset with the data from Marques et al., 2018 leading to ∼4 million bouts. A last reference dataset consisted of prey capture events containing a total of 1700 short capture swims and 3200 long capture swims. All reference datasets were acquired at 700 fps but we used slightly different tracking methods in addition to different area sizes and depths. This variability was critical to ensure the classifier’s ability to tackle out of sample datasets.

We applied the segmentation and classification MATLAB code from Marques et al., 2018 to get the reference bout category. This dataset was split into 80-20% train and test for training the classifier.

For some categories, we applied hierarchical clustering to yield subcategories labels:

- the subcategory for double O-bend was based on Randlett et al., 2019.
- Burst Swim were split since we found different speed tuning within the subcategories (see Fig. S6).
- Short-latency C-start were split based on differences for visual or acoustic stimuli (see Fig. S6).
- The Long capture swims were split according to the sign of the turning tail beat.

Overall, the subcategories lead to a small but significant increase in the mutual information (a statistical measure of dependency between two variables) between subcategories and stimuli than between categories and stimuli (mean = 0.1076 nats vs mean=0.1025 nats, p=4e-9, Wilcoxon signed-rank test, N=108).

#### 3.B. Head-restrained

We collected a dataset from 25 fish performing OMR and Spontaneous Swimming. From this, we extracted 370 5-second epochs containing a variety of tail movements for learning the sparse-coding dictionary.

### Supplementary Note 4: Megabouts freely swimming pipeline

#### 4.A. Data inputs format

All input positions should be provided in millimeters (mm) and angular values in radians. For tail tracking, at least 4 keypoints from the swim bladder to the tail tip are required. The tail posture will be interpolated to 11 equidistant points, thus giving 10 angular values for it. For trajectory tracking, at least two keypoints are necessary to estimate the larva’s position and head direction. In low spatial resolution conditions, we found that tracking two keypoints could be reliably done using SLEAP to estimate the head orientation (see Supp. Note 2.C).

#### 4.B. Smoothing

An option for interpolating the data is available, but it is only applied when a few short segments (∼10 ms) are missing from a high-frequency recording. The head trajectory is smoothed using a 1-euro filter, since it can outperform the Kalman filter while being very simple to adjust on a new dataset (Casiez et al., 2012). An option is provided to denoise the tail angle using a principal component analysis bottleneck and keeping only 4 PCs similarly to Mearns et al., 2020. The tail angle is also smoothed using the Savitzky-Golay filter.

All default parameters expressed in units of time are scaled according to the frame rate. For instance, the window length of the Savitzky-Golay is 15 milliseconds for a second-order polynomial. The corresponding number of frames is computed according to the recording frame rate. This ensures that the parameter tuning is minimal.

#### 4.C. Segmentation

##### 4.C.1. Segmentation from tail angle

Zebrafish larvae swim in a burst-and-glide pattern with discrete bouts lasting typically between 100 and 200 ms. To identify tail bouts, we compute the speed of the tail. Since first-order numerical differentiation typically introduces noise, we instead used smooth noise-robust differentiators (Holoborodko, 2008) that are optimized to produce a smooth estimation of the derivative of a numerical signal. We compute the derivative of the angles along the tail, cumulate the rectified value of the speed and apply a low pass filter. The bouts can then be identified by applying a threshold on the tail angle. See results in Fig. S3i.

##### 4.C.2. Segmentation from the trajectory

We computed the kinematic activity or trajectory speed using angular and positional speed (estimated using Holoborodko, 2008). This measure has the property of peaking near the first tail beat. A peak-finding algorithm is employed to find the approximate location of the first tail beat. The onset and offset are then computed by thresholding at a given percentage relative to the peak (typically 20%). See results in Supplementary Fig. S3ii,iii.

#### 4.D. Swim bout characterization

Following the segmentation, the swim bout trajectory will be centered and aligned according to the fish position and direction at the onset (see Fig. S4a). This will put all swims in a common reference frame. The tail angle input to the transformer is limited to the first 7 out of 10 values since we found no improvement using the segments closer to the tip of the tail, where tracking errors are more common. A tensor containing all swim bouts from a given experiment is fed to the Transformer neural network, which outputs the probability distributions over bout categories and subcategories. Additionally, the network predicts the location and direction of the first tail beat, enabling the calculation of latency to stimulus onset and providing directional information about the swim bout.

#### 4.E. Transformer Neural Network

##### 4.E.1. Data Augmentation

The training dataset described in Supp. Note 3.A consists of tail and trajectory at 700 fps. During training, we used the raw recording values since the downsampling value obtained from smoothing at 700 fps could lead to an artificially high signal-to-noise ratio. Random Gaussian noise was added to 10% of the sample. The head angle was also flipped by 180^*°*^ for random frames to enhance robustness against tracking error. The swim bouts were shifted in time to yield robustness to a difference in onset detection. Down sampling was applied to yield an effective frame rate between 10 and 700 fps (see Fig. S4c). We also randomly masked the tail information in 50% of the cases to get accurate classification from head trajectory only (see Fig. S4c). The trajectory was also removed in 5% of the training examples.

##### 4.E.2. Network Architecture

The network architecture is illustrated in Fig. S4b. The maximum number of tokens was 140, corresponding to 200 ms at 700 fps. The positional encoding was computed as follows:

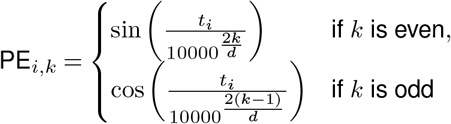

where:

- PE_*i,k*_ represents the positional encoding matrix, where *i* is the position in the sequence, and *k* ∈ {0, 1, 2, …, 63} is the embedding index.
- *d* = 64 is the embedding dimension for tail bouts and position.
- *t*_*i*_ is the timestamp relative to the onset of movement of the corresponding camera frame in milliseconds.

A learned classification token was used at position 0, leading to 141 input tokens. A learned mask was used to fill in missing data for tokens where either tail or trajectory was missing. We used a multi-head self-attention with 8 heads and 3 layers. The output of the network was linearly projected to obtain logits for the subcategories. Let 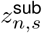 denote the logits for subcategory *s* and sample *n*, and let 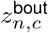 represent the logits for bout category *c* and sample *n*. The logits for the bout categories were derived from the subcategory logits using the following formula:

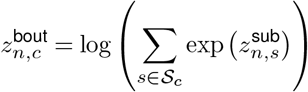

where:

- *n* indexes the samples in the batch.
- *s* indexes the subcategories.
- *c* indexes the bout categories.
- *𝒮* _*c*_ is the set of subcategories that belong to bout category *c*.

Here, 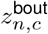 represents the logit for bout category *c* for sample *n*, and the summation is performed over the subcategory indices *s* that belong to bout category *c*.

Another linear projection from the output classification token was used to predict the direction of the first half-beat. A sigmoid activation function was applied to predict the first tail beat location relative to the first frame.

The network was implemented in PyTorch.

##### 4.E.3. Network Training

Random data augmentation was applied at the beginning of each epoch. For each batch, the input data were balanced across categories. The loss was defined as the weighted sum of the cross entropy value for the bout category, subcategory and sign and the mean squared error for the first half beat location. The weights were computed to give roughly similar value to the four loss terms at the network initialization. We used an Adam optimizer with a learning rate of 1e-4, a batch size of 50 swim bouts, and a dropout rate of 0.1. It ran until convergence on the training set.

#### 4.F. Pipeline evalutation

##### 4.F.1. Segmentation evaluation

Fig. S3c shows the accuracy of segmentation based on trajectory compared with segmentation based on tail speed. This metric quantifies how often the onset of movements detected by each segmentation method occurs within a specified time window of each other. To calculate this index, we started by identifying the time window within which we consider two movement onsets to be matching (here, we used 75ms, which corresponds to one and a half samples at 20fps). Then, we examined each onset from the first segmentation method to see if there is at least one onset from the second method occurring within this window, and we did the same in reverse. The coincidence index is then determined by counting these matches and comparing the count to the total number of movement onsets detected by both methods. This index provides a straightforward way to assess the degree of synchrony between two segmentation methods, with higher values indicating greater alignment in detecting movement onsets.

##### 4.F.2. Transformer evaluation

To evaluate the classifier’s performance, we masked and downsampled swim bouts in the test set to simulate different frame rate and tracking configurations. To ensure the performance was evaluated with imperfect bout alignment, bouts were segmented with random shift, yielding the first tail beat to be positioned with equal probability between -14ms and +70ms relative to the first token. This distribution was chosen to roughly match the support of the distribution of segmentation error at 20 fps (see Fig. S3b).

Since the distribution of bouts strongly depends on the sensory context, we computed the balanced accuracy of bout category and sub-category to evaluate the network performance at different fps and with and without tail tracking (Fig. 1c and Fig. S5b). We found that the balanced accuracy for ‘tail + trajectory tracking’ at 700 fps was 89%. It should be noted that we should not expect 100% accuracy. Indeed, the bout categories are defined according to the unsupervised clustering of Marques et al., 2018. This classification relies on the estimation of a large number of kinematic parameters extracted for each tail beat within a bout, these measures are very sensitive to shift compared with our strategy. Moreover, the unsupervised clustering of swim bouts corresponds to density peaks in the space of swim bouts parameters (Marques and Orger, 2019) rather than on isolated clusters. Thus, the continuity in kinematic space makes it difficult to separate similar categories, such as Approach Swim and Slow1, as can be seen from the confusion matrix in Fig. S5a.

A critical metric for ensuring the quality of the classification is to check that the bout category is informative about the fish’s sensory experience. As reported in the main text, the mutual information was higher using the category from Megabouts than with the MATLAB version (mean = 0.1025 nats vs mean=0.0934 nats, p=4e-17, Wilcoxon signed-rank test, N=108). Along similar lines, another criteria for assessing movement clustering is that the information should be predictive of future actions (Berman et al., 2016). We found that the categories from Megabouts were more predictive of future movements during spontaneous activity (for future movement, we used the category defined according to the MATLAB reference code). (mean = 0.1209 nats vs mean=0.1197 nats, p=3e-6, Wilcoxon signed-rank test, N=108). We hypothesized that this improvement results from the augmentation strategy leading to a more robust estimation of bout categories.

In the Zebrabox assay we are missing a high-speed reference evaluation of the bout categories. To overcome this drawback, we used the fact that fish generate O-Bends with high probability when the light is switched off (Burgess and Granato, 2007; Randlett et al., 2019). Indeed, we saw that for all the well sizes, a peak in O-Bend probability is present after the light is switched off (see Fig. S7).

### Supplementary Note 5: Megabouts head-restrained pipeline

#### 5.A. Preprocessing and detrending

The first preprocessing steps are similar to Supp. Note 4.B. The baseline removal is an additional step for pre-processing head-restrained tracking. In head-restrained conditions, tail angles commonly exhibit a slow varying baseline. This low-frequency component is first subtracted before running the sparse coding algorithm. The algorithm used to fit the baseline is obtained from the python library pybaselines (Erb). This slow baseline could be relevant and has been associated with activity in the nucleus of the medial longitudinal fascicle (nMLF) (Dal Maschio et al., 2017).

#### 5.B. Sparse coding

Convolutional sparse coding is an unsupervised method that decomposes a continuous image or multidimensional time series into the sparse contributions of motifs. The ensemble of motifs forms a dictionary that is learned from the data to reconstruct the original signal from a convolution of motifs with a sparse code (see Fig. S8). The specific algorithm we used was designed to recover the individual keystrokes from piano auditory recordings (Wohlberg, 2017; Cogliati et al., 2017). We collected data on the tail angle of head-restrained zebrafish and used the algorithm to learn a dictionary of 4 motifs: slow forward movements, fast forward movements, turns, and escape/struggle-like movements. We found that although the duration of tail movements could exceed 1s in head-fixed conditions, the duration of motifs did not exceed 200ms, suggesting that the basic time scale is similar to freely swimming bouts. From the sparse code, the tail angle time series can be decomposed into the contributions of the four motifs.

#### 5.C. Sparse coding dictionary learning

The sparse deconvolution was computed using the *ConvBPDN* algorithm from SPORCO. The dictionary learning used the *ConvBPDNDictLearnConsensus* method. To have dictionary motifs with matching first half beat locations we employed a learning schedule. For each iteration, we increased the sparsity and shifted the learned motif so that their first half beat would be localized at the beginning. After 10 iterations the algorithm converged.

## Supplementary Videos

**Figure SV1.**
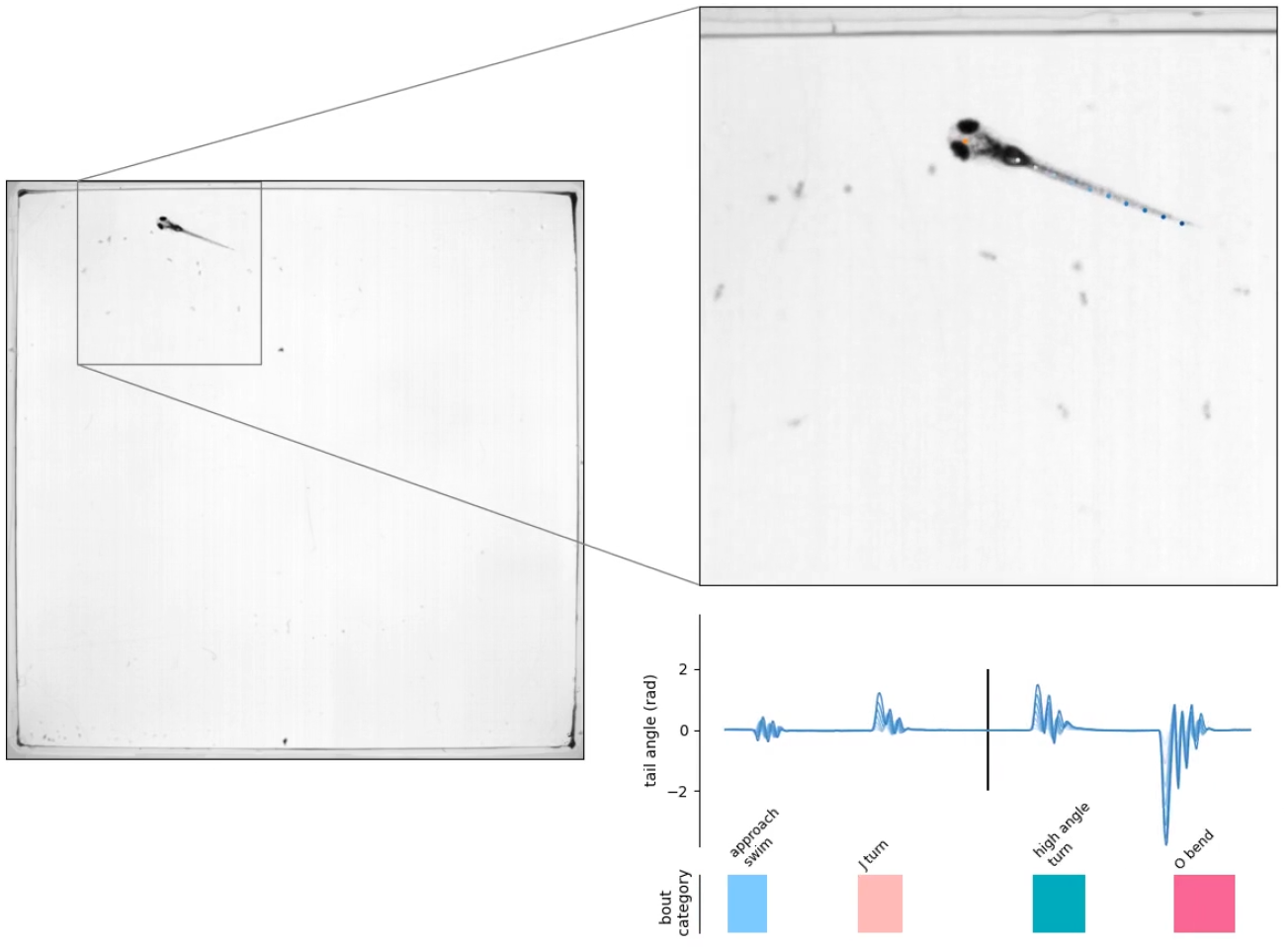
Use case of Megabouts for computing ethogram from ‘tail+trajectory tracking’ at 700 fps.

**Figure SV2.**
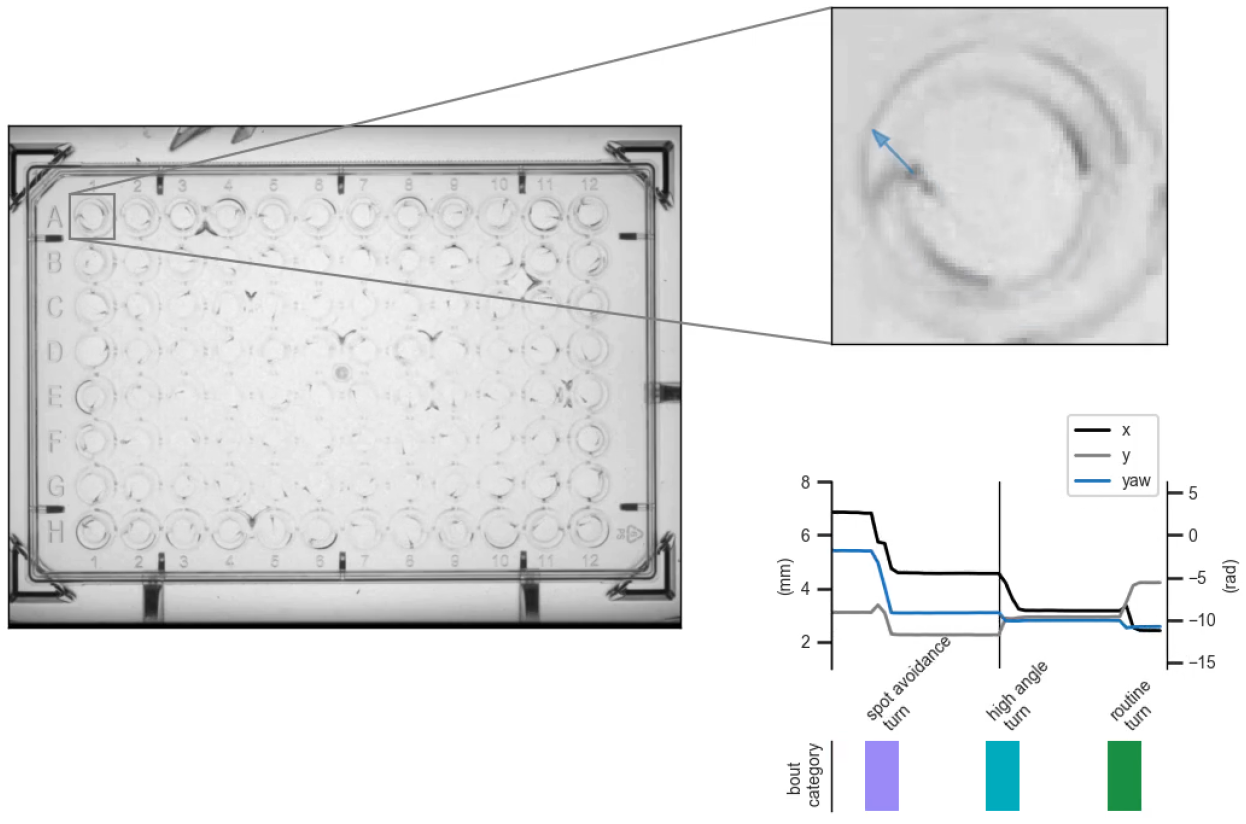
Use case of Megabouts for computing ethogram from ‘trajectory tracking’ at 25 fps.

**Figure SV3.**
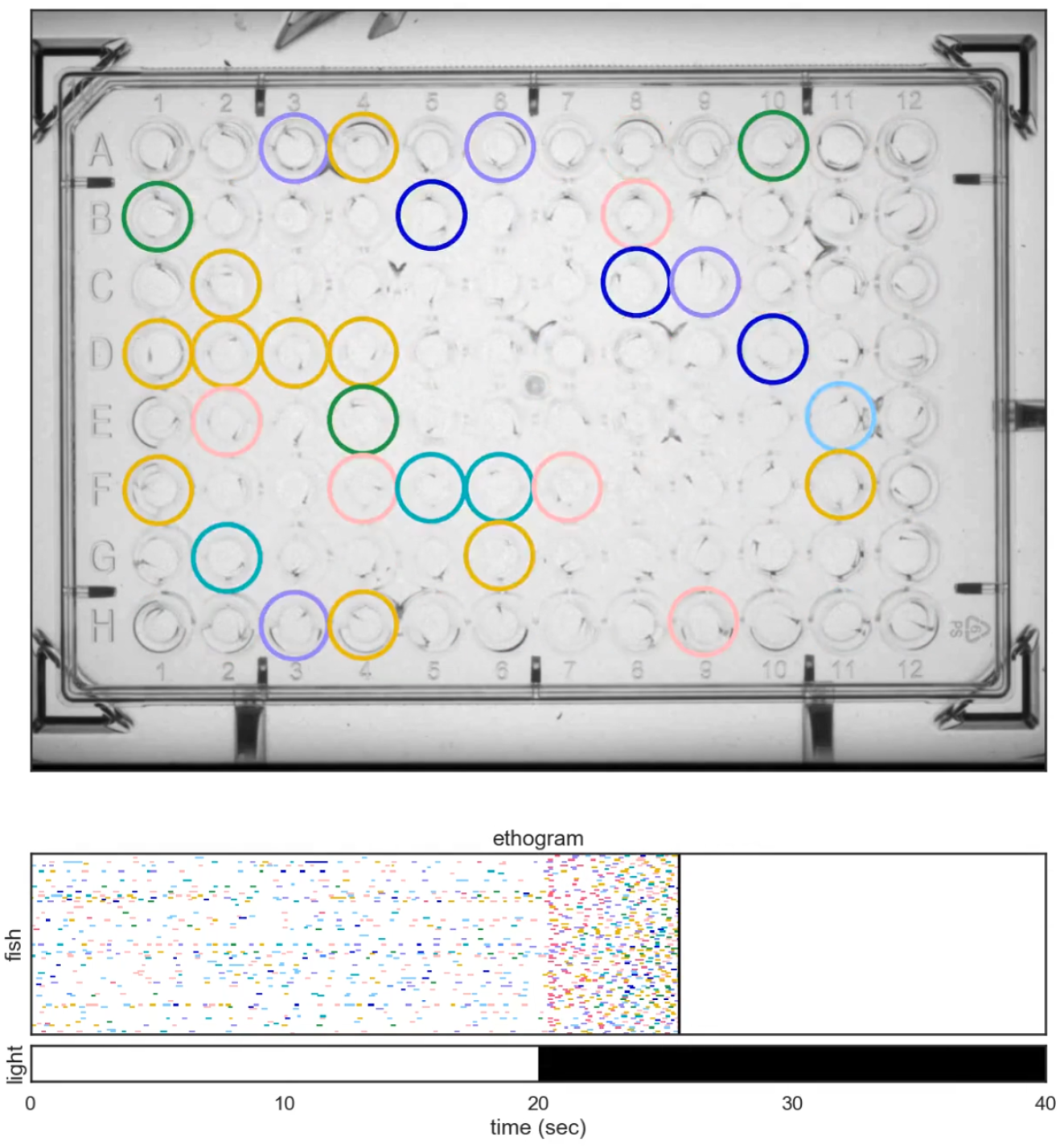
High-throughput recording of ethogram during light on/off stimuli.

